# The first chromosomal-level genome assembly of an allotetraploid oyster

**DOI:** 10.1101/2025.02.25.639704

**Authors:** Ao Li, Jinlong Zhao, Mingjie Zhao, Mengshi Zhang, Meitong Huo, Jinhe Deng, Luping Wang, Wei Wang, Haigang Qi, Yalin Li, Xiaoyu Li, Jie Fu, Xirui Guo, Zhe Xu, Li Li, Ximing Guo, Guofan Zhang

**Affiliations:** Key Laboratory of Breeding Biotechnology and Sustainable Aquaculture (CAS), Institute of Oceanology, Chinese Academy of Sciences, Qingdao, China; Laboratory for Marine Biology and Biotechnology, Qingdao Marine Science and Technology Center, Qingdao 266237, China; Shandong Province Key Laboratory of Experimental Marine Biology, Institute of Oceanology, Chinese Academy of Sciences, Qingdao 266071, China; University of Chinese Academy of Sciences, Beijing 100049, China; National and Local Joint Engineering Laboratory of Ecological Mariculture, Qingdao 266071, China; Oyster Industrial Technology Institute of Zhanjiang, Southern Marine Science and Engineering Guangdong Laboratory (Zhanjiang), Zhanjiang 524000, China; Shandong Center of Technology Innovation for Oyster Seed Industry, Qingdao 266000, China; Qingdao Frontier Ocean Seed Company Ltd., Qingdao 266105, China; Haskin Shellfish Research Laboratory, Department of Marine and Coastal Sciences, Rutgers University, Port Norris, NJ 08349, USA

## Abstract

Tetraploid oysters are used to cross with diploids to produce triploid oysters that have become an important part of the oyster aquaculture industry worldwide. Although most tetraploid oysters are artificially induced autotetraploids, allotetraploids can be produced between closely related species, providing new opportunities for polyploid breeding and studying genome interactions. Using PacBio sequencing, Illumina sequencing, and high-throughput chromosome conformation capture scaffolding, we assembled the first high-quality genome assembly of an artificially induced allotetraploid between the Pacific oyster *Crassostrea gigas* and Portuguese oyster *Crassostrea angulata*. The assembled genome is 1.23 Gb, with a contig N50 of 2.56 Mb and a scaffold N50 of 57.22 Mb, and anchored to 20 chromosomes. The assembly contains 58,330 protein-coding genes, 98.34% of which are functionally annotated. The heterozygosity and the ratio of repetitive sequences is 5.50% and 46.43%, respectively. This chromosomal-level genome assembly of an allotetraploid oyster provides a valuable genetic resource for studying genome biology of molluscs and for advanced breeding of polyploids that are critical for the oyster aquaculture industry.

## Background & Summary

Oysters are among the most important aquaculture species worldwide, accounting for an annual production of ∼7 million metric tons (World Food and Agriculture – Statistical Yearbook 2024 (fao.org)). Genetic improvement including selective breeding, hybridization and polyploidization, plays an important role in supporting oyster aquaculture^1^. One of the most significant advances over the last four decades is the development of triploid oysters. Because of their superior growth, improved meat quality and sterility, triploids have become one of the most popular stocks for oyster aquaculture^1–3^. The commercialization of triploid oysters has contributed significantly to oyster aquaculture, especially for the Pacific oyster *Crassostrea gigas* and Eastern oyster *Crassostrea virginica*), in meeting the market demand around the world^4,5^. Triploid oysters now account for 30-70% of the cultured oysters in major producing countries such as France, Australia, USA and China (Guo 2021). Originally, triploids were induced by retaining the second polar body in newly fertilized eggs with chemicals such as cytochalasin B (CB) or 6-dimethylaminopurine (6-DMAP)^6,7^. However, chemical induction had low efficiency which hindered commercial production. The successful development of tetraploid oysters by Guo and Allen^8^ (1994) made it possible to produce mated triploids by mating diploids x tetraploids, which is 100% effective without any use of toxic chemicals^5^. Nowadays, triploid oysters are commercially produced through diploid x tetraploid crosses. Thus, successful production and breeding of tetraploids are critical for oyster aquaculture that is heavily dependent on triploids.

The Guo and Allen method for tetraploid induction involves blocking the release of polar body I in eggs from triploid Pacific oysters fertilized by haploid sperm (3n ♀ × 2n♂), which successfully introduced the first autotetraploid Pacific oyster^8,9^. While it is challenging and difficult to reproduce, tetraploids can also be obtained using normal diploid eggs (2n × 2n)^10^. Induction of tetraploidy has also been reported in several oyster species including *C. gigas, C. virginica*, *C. angulata*, *Crassostrea hongkongensis, Crassostrea sikamea*, and tropical oysters *Crassostrea belcheri* (Sowerby) and *Crassostrea iredalei* (Faustino)^11–14^, although it is not clear whether breeding populations of tetraploids have been established the latter two species. Most of the tetraploid oysters produced so far were autotetraploids. Allotetraploids can also be produced between species that can hybridize. Tetraploid genomes represent a new state of whole genome duplication that may be instable and go through rapid reorganization and evolution. With two different genomes, allotetraploids may be more stable and provide a rare opportunity for studying genome interaction. They may also generate new genotypes by combining characteristics of two species and produce superior triploids for aquaculture. High-quality assemblies of diploid genomes have been produced for oyster species and led to advances in oyster biology and adaptation^15–22^. The sequencing and analyses of tetraploid genomes may provide insights into the biology and evolutionary potential of tetraploids.

We previously produced allotetraploid oysters between the Pacific oyster *C. gigas* and Portuguese oyster *C. angulata*, two closely related species that dominate oyster aquaculture production^23^. In this study, we used long reads generated by PacBio sequencing, short reads generated by Illumina sequencing, and high-throughput chromosomal conformation capture (Hi-C) analysis and constructed the first high-quality chromosomal-level genome assembly of the allotetraploid oyster. The final genome size is 1,230.39 Mb in 717 contigs, with a contig N50 length of 2.56 Mb and a scaffold N50 length of 57.22 Mb. More than 90% of contigs (1,108.13 Mb) were anchored on 20 chromosomes. The assembly contains 571.24 Mb (46.43%) of repetitive sequences and 7,961 noncoding RNAs. Using *de novo* prediction, mRNA transcripts and homolog-based strategies, a total of 58,330 protein-coding genes were predicted, and 98.34% of which (57,360) were annotated in the publicly available NCBI RefSeq non-redundant protein, eggNOG, KEGG, SWISS-PROT, Pfam, TrEMBL, GO, and KOG databases. This allotetraploid oyster genome assembly provides a valuable resource for studying genome interaction and evolution, polyploid biology and genetic improvement of polyploid oysters for aquaculture.

## Methods

### Sample and sequencing

The allotetraploid oyster was artificially induced between the Portuguese oyster *C. angulata* and the Pacific oyster *C. gigas*. First, allotriploids were produced in 2015 by mating diploid *C. angulata* and autotetraploid *C. gigas.* Second, allotetraploids were produced in 2018 with the Guo and Allen method^8^ using eggs from the allotriploids and sperm from diploid *C. angulata*. Subsequently, allotetraploids were reproduced by 4n x 4n crosses for three generations. For this study, one allotetraploid oyster was sampled on 05/27/2024 from the F_3_ allotetraploids that were produced in 2022. The tetraploidy of the sampled oyster was confirmed by flow cytometry. Adductor muscle was collected and flash-frozen in liquid nitrogen, and then used for genomic DNA extraction (with ∼30 mg tissue) using the DNeasy Blood & Tissue Kit (Qiagen, Hilden, Germany). Agarose (1.0%) gel electrophoresis, Qubit (Invitrogen, Qubit^TM^3Flurometer) and NanoDrop 2000 spectrophotometer (Thermo Fisher Scientific, Waltham, MA, USA) were used to determine DNA concentration and quality. The genomic DNA was used to build sequencing libraries, including 15-kb insert PacBio HiFi library and 150-bp insert Illumina paired-end library.

High-molecular weight (HMW) gDNA was prepared for PacBio HiFi read production and libraries was constructed using the PacBio Template Prep Kit 1.0 according to standard protocol of Template Preparation using BluePippin size selection (Pacific Biosciences, USA). Sequencing of genomic libraries was performed on two cells using the self-testing high-precision CCS mode on the PacBio Sequel II system. A total of 34.37 Gb of HiFi long-read data with a read N50 length of 16.59 Kb (average read length of 16.18 Kb) was obtained, resulting in 27.94-fold coverage of the allotetraploid oyster genome size.

The short-insert library was constructed using the NR604-VAHTS Universal V6 RNA-seq Library Prep Kit (Vazyme), and then was sequenced by the Illumina NovaSeq 6000 platform using the paired-end model (PE 150) following the standard protocol (Illumina Inc., San Diego, CA, USA). A total of 156.82 Gb (127.50-fold coverage) of clear reads with a Q30 of 93.52% were obtained to assess allotetraploid oyster genome size.

The Hi-C libraries were also constructed for genome assembly^24,25^. The same fresh adductor muscle was crosslinked with 1.0% formaldehyde and then was terminated with 0.2 M glycine. Libraries were generated according to the manufacturer’s instructions: 1) digestion with *HindIII* restriction enzyme, 2) labeling using Biotin-14-dATP (Thermo Fisher Scientific, USA), 3) ligation with T4 DNA ligase, 4) physically shearing into 300-700 bp fragments, 5) selectively capture using streptavidin magnetic beads. Illumina HiSeq 6000 platform was used for sequencing. We obtained 147.05 Gb (119.55-fold coverage) of clean data.

For genome annotation, we collected tissues from four organs (gill, mantle, adductor muscle and labial palp) for RNA-seq. Total RNA was extracted from each of tissues and then were equally mixed into 1 sample. The RNA mixture was used for library construction and sequencing by the Illumina NovaSeq 6000 platform following the standard protocol (Illumina Inc., San Diego, CA, USA). A total of 7.17 Gb of clear data was yield.

### Genome assessment and assembly

Illumina paired-end clear reads (156.82 Gb) was used to survey genome feature of allotetraploid oyster via the *k-mer* method. GenomeScope v2.00^26^ (parameters: -k 19 -p 4 -m 1000000000) and Jellyfish v2.1.4^27^ (parameter: -h 1000000000) were used for *k-mer* count histogram (*k* = 19) (Fig. 1). Estimation of genome size followed the formular of G = N *_k-mer_*/Daverage *_k-mer_*, where N *_k-mer_* is the total number of *k-mers*, Daverage *_k-mer_* is the average depth of *k-mer*s, G is genome size. The survey results showed that the haploid genome size of allotetraploid oyster was estimated to be 544.56 Mb with the heterozygosity, repetitive sequence ratio and GC content of 5.50%, 46.69% and 34.61%, respectively (Table 1). In total, 34.37 Gb of HiFi long-reads were used for assembly using the Hifiasm v0.19 software^28^ with default parameters, resulting in a total length of 1,815.36 Mb comprising 1,610 contigs with a contig N50 length of 2.29 Mb for the allotetraploid oyster (Table 2).

**Fig. 1.**
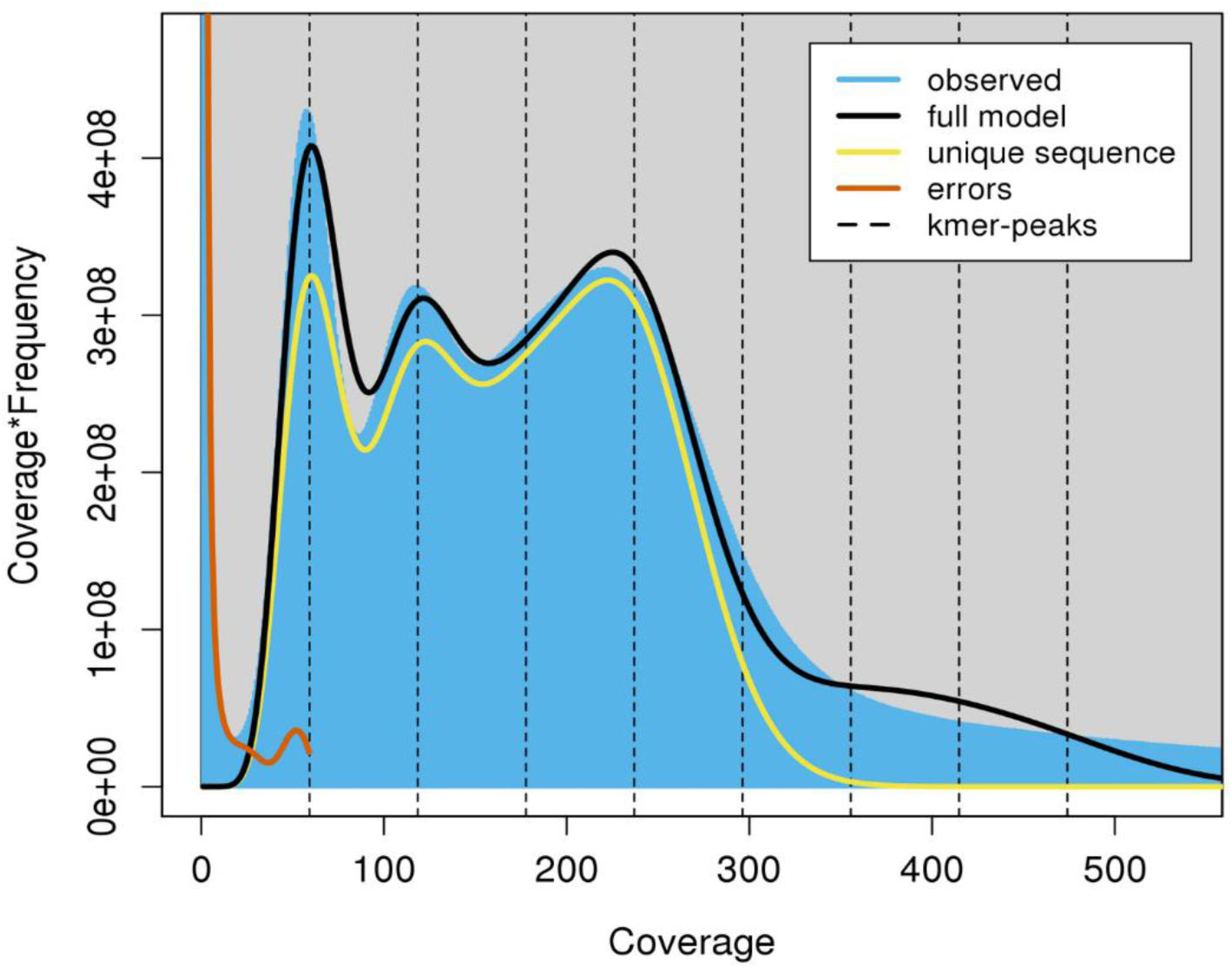
The 19-mer frequency distribution in the allotetraploid oyster genome. The percentages of different genotypes are: aaaa [94.5%], aaab [3.15%], aabb[1.99%], aabc [0.001%] and abcd [0.361%]. The X-axis is the k-mer coverage, and Y-axis represents the product of frequency by coverage of the k-mer.

**Table 1.** Characteristics of the allotetraploid oyster genome based on *k-mer* analysis.

| $K_{mer}$ | Depth | Genome size (Mb) | Heterozygous rate (%) | Repeat rate (%) | GC content (%) |
| --- | --- | --- | --- | --- | --- |
| 19 | 143.99× | 1089.12 | 5.50 | 46.69 | 34.61 |

**Table 2.**
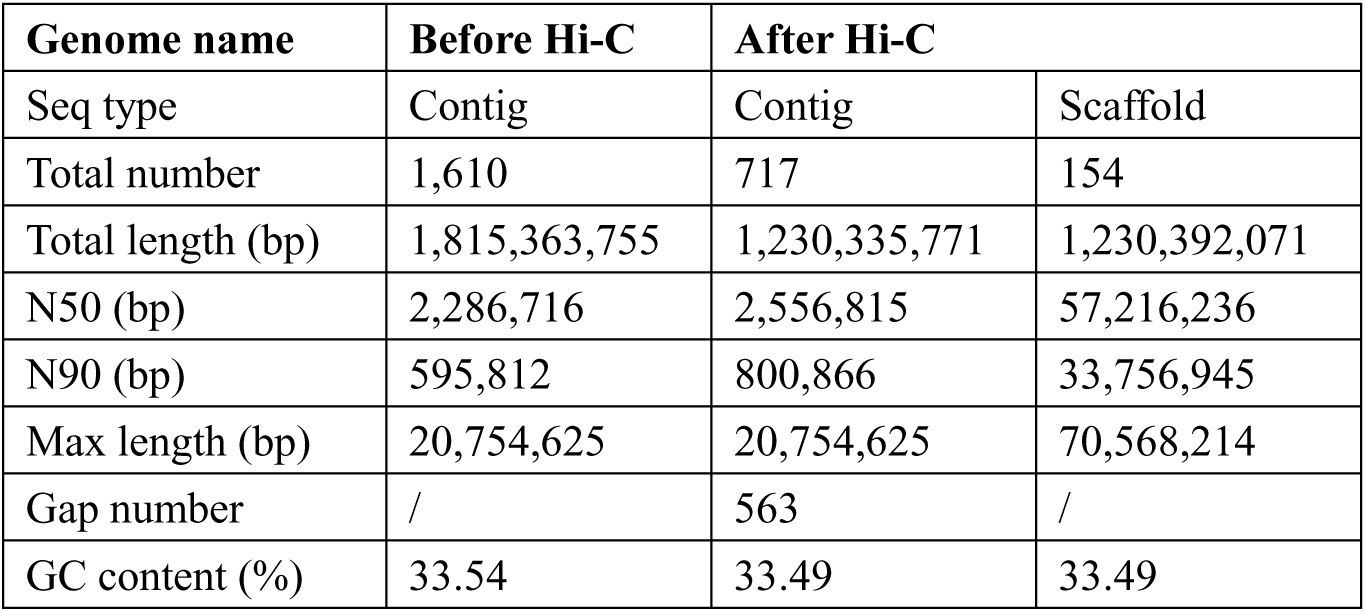
Assembly statistics of the allotetraploid oyster genome.

### Chromosomal-level assemble with Hi-C

For anchored contigs, 491,784,314 clean reads generated from the Hi-C data were mapped to the assembly using BWA v0.7.17-r1188^29^ with default parameters. Valid interaction pairs (125,471,951 pairs) were defined as paired reads with mate mapped to a different contig and then were used to do the Hi-C associated scaffolding using HiC-Pro v2.10.0^30^ that can also filter invalid interaction pairs including self-ligation, non-ligation, PCR amplification, random break, extreme fragments and so on. The LACHESIS v2.0.1^31^ was used for agglomerative hierarchical clustering, sorting and orientation (cluster_min_re_ sites = 544; cluster_max_link_density = 2; order_min_n _res_in_trunk = 908; order_min_n_res_in_shreds = 870). All of 1,610 contigs were clustered into 717 groups (contigs after Hi-C) with a contig N50 length of 2.56 Mb and scaffold N50 length of 57.22 Mb (Table 2), and 91.35% (655) was anchored on 20 chromosomes. Finally, 583 contigs were successfully sorted and oriented with the total length of 1,108.13 Mb for allotetraploid oyster (Table 3). Chromatin contact matrix was built by Juicebox v1.5^32^, and the 20 chromosomes show clearly distribution in the heatmap, with distinct interaction signal around the diagonal within chromosome and between adjacent chromosomes (Fig. 2). Moreover, we carried out collinearity analysis of the assembled allotetraploid genome with the original diploid *C. gigas* and *C. angulata* reference genome using Diamond v0.9.29.130^33^ (e<1e−5, C score>0.5) and MCScanX^34^ (MCScanX -s 5 -m 5). The pronounced co-linearity relationships indicated highly conserved gene blocks among allotetraploid oyster, diploid *C. gigas* and *C. angulata* (Fig. 3).

**Table 3.** Statistics of allotetraploid oyster genome sequence length (chromosome level).

| Group | Ordered number | Length (bp) | Percentage |
| --- | --- | --- | --- |
| Chr01A | 46 | 70,563,714 | 6.10 |
| Chr01B | 27 | 68,445,240 | 5.92 |
| Chr02A | 22 | 59,591,169 | 5.15 |
| Chr02B | 37 | 56,902,171 | 4.92 |
| Chr03A | 26 | 61,522,249 | 5.32 |
| Chr03B | 19 | 54,652,417 | 4.73 |
| Chr04A | 42 | 58,409,816 | 5.05 |
| Chr04B | 36 | 51,572,103 | 4.46 |
| Chr05A | 39 | 55,458,002 | 4.80 |
| Chr05B | 21 | 51,760,265 | 4.48 |
| Chr06A | 28 | 57,213,536 | 4.95 |
| Chr06B | 23 | 55,030,019 | 4.76 |
| Chr07A | 28 | 60,712,823 | 5.25 |
| Chr07B | 25 | 59,918,863 | 5.18 |
| Chr08A | 27 | 52,204,945 | 4.52 |
| Chr08B | 28 | 43,426,356 | 3.76 |
| Chr09A | 31 | 63,045,174 | 5.45 |
| Chr09B | 41 | 59,112,309 | 5.11 |
| Chr10A | 14 | 34,837,826 | 3.01 |
| Chr10B | 23 | 33,754,745 | 2.92 |
| Total | 583 | 1,108,133,742 | 95.84 |
| Unplaced | 72 | 48,078,193 | 4.16 |

**Fig 2.**
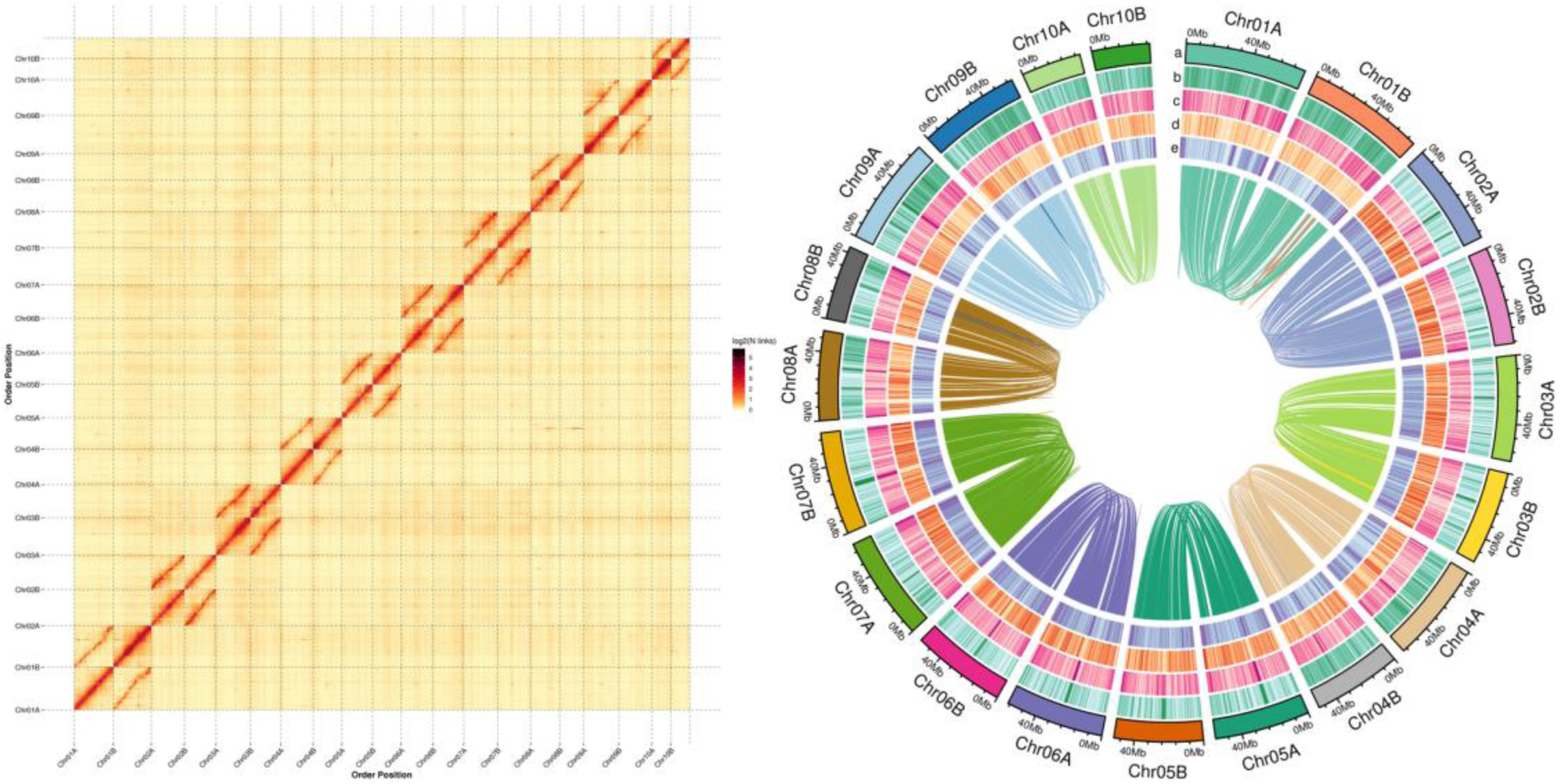
Genome assembly of the allotetraploid oyster. (**a**) Hi-C assembly of chromosome interactive heat map. Color block indicates intensity of interaction from yellow (low) to dark-red (high). (**b**) Circos plot of the assembly. The a, b, c, d and e indicate chromosome ideograms, TE density, SSR density, gene density and GC content.

**Fig 3.**
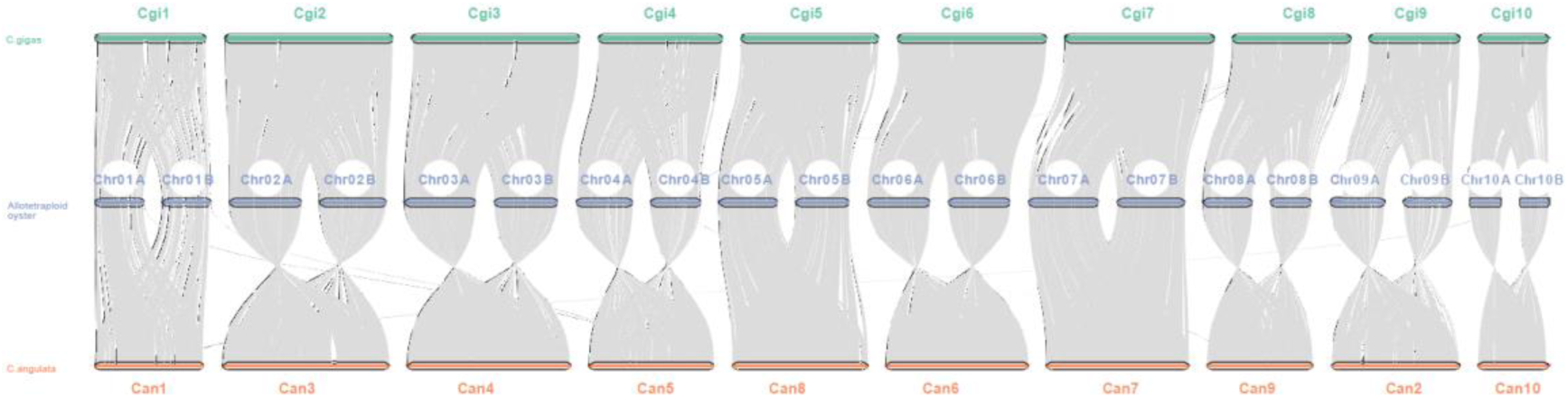
Collinearity analysis among genomes of the allotetraploid oyster, diploid *C. gigas* and *C. angulata*.

### Repeat sequences annotation

Whole-genome repeat sequences, including tandem repeats and transposable elements (TEs), were annotated using the combined strategy of *ab initio* prediction and homology alignment. The MIcroSAtellite identification tool (MISA v2.1^35^) and Tandem Repeat Finder (TRF v409^36^, 2 7 7 80 10 50 500 -d -h) were used to predict tandem repeats, which yielded a total of 83.22 Mb of tandem repeats (6.76% of the genome assembly) (Table 4). For TEs, a customized repeat library was built using RepeatModeler v2.0.1^37^ (BuildDatabase -name && RepeatModeler -pa 12), which can initiate two *de novo* repeat finding programs of RECON v1.0.8^38^ and RepeatScout v1.0.6^39^. The library was then classified by RepeatClassifier with default parameters according to the public databases of Dfam v3.5^40^ and Repbase v19.06^41^. The LTRharvest v1.5.10^42^ and LTR_finder v2.8^43^ (ltr_finder -w 2 -C -D) were used to identify full-length long terminal repeat retrotransposons (fl-LTR-RTs). High-quality intact fl-LTR-RTs and non-redundant LTR library were then generated by LTR_retriever v2.9.0^44^. We combined the above *de novo* TE sequences libraries with public databases to construct non-redundant species-specific TE library, which was then used to identify and classify the final TE sequences using homology search of RepeatMasker v4.1.2^45^ (repeatmasker -nolow -no_is -norna -engine wublast -parallel 8 -qq). A total of 488.03 Mb of TEs were identified, accounting for 39.67% of genome assembly. Among TEs, DNA transposons and retroelements accounted for 26.85% (330.34 Mb) and 12.82% (157.68 Mb) of the genome assembly, respectively (Table 4).

**Table 4.** Annotation of repeat sequences for the assemble allotetraploid oyster genome. Note: Number: the number of repetitive sequences; Length: the total length of predicted repetitive sequences; % in Genome: the proportion of repetitive sequences in the total genome.

| Class | Type | Number | Length (bp) | % in Genome |
| --- | --- | --- | --- | --- |
| Transposable elements | DIRS | 216 | 65,777 | 0.0054 |
|  | LINE | 89,279 | 28,303,157 | 2.30 |
|  | LTR | 391,338 | 128,886,291 | 10.48 |
|  | SINE | 4,242 | 423,085 | 0.034 |
|  | DNA | 1,029,960 | 330,341,186 | 26.85 |
|  | srpRNA | 1 | 289 | 0.000023 |
|  | Unknown | 123 | 12,545 | 0.0010 |
|  | Total | 1,515,159 | 488,032,330 | 39.67 |
| Tandem repeats | microsatellite (1-9 bp units) | 644,644 | 13,711,047 | 1.11 |
|  | minisatellite (10-99 bp units) | 205,033 | 25,830,275 | 2.10 |
| | satellite ($\geq 100$ bp units) | 50,088 | 43,674,118 | 3.55 |
|  | Total | 899,765 | 83,215,440 | 6.76 |

### Noncoding RNAs and pseudogene annotation

For noncoding RNAs annotation, miRNA, rRNA, tRNA, snoRNA and snRNA were identified by specific approaches. The miRNA was identified against miRBase database^46^. Based on the Rfam v14.5^47^ database, rRNA and tRNA were identified by tRNAscan-SE v1.3.1^48^ and barrnap v0.9^49^ (barrnap --kingdom euk --threads 1) respectively, and snoRNA and snRNA were identified by Infernal v1.1^50^ (cmscan --cpu 3 --rfam). In total, 7,710 tRNA, 179 rRNA and 72 miRNA were predicted (Table 5).

**Table 5.** Noncoding RNAs and pseudogene annotation of the assemble allotetraploid oyster genome.

| RNA classification | Number | Pseudogene | Value |
| --- | --- | --- | --- |
| miRNA | 72 | Number | 362 |
| rRNA | 179 | Total length | 2,043,162 |
| tRNA | 7,710 | Average length | 5.64 Kb |

The GenBlastA v1.0.4^51^ and GeneWise v2.4.1^52^ were used to identify homologous pseudogenes after excluding functional genes (genblasta -P wublast -pg tblastn) and to search for immature stop codon and frameshift mutations (genewise -both -pseudo), respectively. We obtained 362 pseudogenes with average length of 5.64 Kb (Table 5).

### Protein-coding gene prediction and functional annotation

Three approaches, *de novo* prediction, homology-based prediction, and mRNA-based prediction, were applied for protein-coding gene prediction in the allotetraploid genome. Two *ab initio* gene-prediction software, Augustus v3.1.0^53^ and SNAP v2006-07-28^54^, were used for *de novo* gene models prediction in the repeat-masked assembly (hard-masking). For homology-based prediction, protein sequences of four well-annotated species (*C. gigas*, *C. angulata*, *C. ariakensis* and *Danio rerio*) were downloaded and aligned to the repeat-masked genome assembly. Then, the GeMoMa v1.7^55^ (run.sh mmseqs) was used to predict gene model based on sequence alignment. The 7.17 Gb clean data from RNA-seq was used for mRNA-based prediction. The Hisat v2.1.0^56^ (hisat2 --dta -p 10) and StringTie v2.1.4^57^ (stringtie -p 2) were used to assemble transcripts. The GeneMarkS-T v5.1^58^ was used to predict genes based on transcripts. Finally, the EVidenceModeler (EVM) v1.1.1^59^ was used to integrate all gene models predicted by the above methods, which was then modified by PASA v2.4.1^60^ to generate a weighted and non-redundant gene set. A total of 58,330 protein-coding genes (Table 6) were predicted with average exon number of 7.84 per gene and average gene length of 8.27 Kb (Table 7).

**Table 6.** Gene prediction of the allotetraploid oyster using three methods.

| Method | Gene set | Species | Number |
| --- | --- | --- | --- |
| Ab initio | Augustus | - | 50,539 |
|  | SNAP | - | 85,354 |
| Homology-based | GeMoMa | *C. angulata* | 63,485 |
|  |  | *C. ariakensis* | 62,945 |
|  |  | *C. virginica* | 51,353 |
|  |  | *D. rerio* | 20,585 |
| RNAseq | GeneMarkS-T | - | 45,019 |
|  | PASA | - | 22,259 |
| Integration | EVM | - | 58,330 |

**Table 7.** The comparison of gene models predicted from the allotetraploid oyster, Pacific oyster (*C. gigas*) and Portuguese oyster (*C. angulata*).

| Statistics | Allotetraploid oyster | *C. gigas* | *C. angulata* |
| --- | --- | --- | --- |
| Gene Num | 58,330 | 25,923 | 28,211 |
| Gene Len | 482,454,743 | 330,861,862 | 209,012,942 |
| Ave Gen Len | 8,271 | 12,763 | 7,409 |
| Exon Len | 109,046,919 | 44,765,190 | 42,907,275 |
| Ave Exon Len | 1,869 | 1,727 | 1,521 |
| Exon Num | 457,043 | 218,906 | 217,883 |
| Ave Exon Num | 7.84 | 8.44 | 7.72 |
| CDS Len | 95,082,579 | 44,765,190 | 42,907,275 |
| Ave CDS Len | 1,630 | 1,727 | 1,521 |
| CDS Num | 450,163 | 218,906 | 217,883 |
| Ave CDS Num | 7.72 | 8.44 | 7.72 |
| Intron Len | 373,407,824 | 286,096,672 | 166,105,667 |
| Ave Intron Len | 6,402 | 11,036 | 5,888 |
| Intron Num | 398,713 | 192,983 | 189,672 |
| Ave Intron Num | 6.84 | 7.44 | 6.72 |

For functional annotation of gene models, we searched against public biological functional databases, including Non-Redundant (NR), Evolutionary Genealogy of Genes: Non-supervised Orthologous Groups (eggNOG)^61^, Gene Ontology (GO), TrEMBL, Gene Ontology (GO), EuKaryotic Orthologous Groups (KOG), Kyoto Encyclopedia of Genes and Genomes (KEGG)^62^, SWISS-PROT^63^ and Pfam^64^, using Diamond blastp (Diamond v0.9.29.130^33^, diamond blastp --masking 0 -e 0.001). A total of 57,360 genes (98.34% of the total predicted genes) were functionally annotated (Table 8).

**Table 8.** Statistics of gene functional annotation of the allotetraploid oyster genome assembly.

| Annotation database | Annotated number | Percentage (%) |
| --- | --- | --- |
| GO | 44,241 | 75.85 |
| KEGG | 42,834 | 73.43 |
| KOG | 30,726 | 52.68 |
| Pfam | 45,647 | 78.26 |
| Swissprot | 28,723 | 49.24 |
| TrEMBL | 57,144 | 97.97 |
| eggNOG | 37,488 | 64.27 |
| NR | 56,480 | 96.83 |
| All_Annotated | 57,360 | 98.34 |

### Data Records

The raw PacBio, Hi-C, and Illumina sequencing data are deposited in the NCBI Sequence Read Archive database under the accession numbers: SRR32459897, SRR32456008 and SRR32455876. Moreover, the genomic annotation results have been deposited in the *figshare* database^65^.

### Technical Validation

Four methods were used to evaluate the genome assembly: the mapping of Illumina reads, PacBio HiFi reads, BUSCO assessment, and core gene integrity. The Illumina short-reads and PacBio HiFi-reads were mapped to the assembly using BWA v0.7.17-r1188^29^ and Minimap2 v2.28^66^ to assess the quality, respectively. As shown in Table 9, 99.09% and 99.95% of short-reads and HiFi-reads were mapped to the allotetraploid genome, respectively. The completeness of the assembly were evaluated by Core Eukaryotic Genes Mapping Approach (CEGMA) v2.5^67^ database and Benchmarking Universal Single-Copy Orthologs (BUSCO) v2.0^68^ against the metazoa_odb10. A total of 447 (97.60%) out of 458 conserved eukaryotic core genes from the CEGMA database and 938 (98.32%) out of the complete 954 BUSCO orthologous groups were identified in the assembled genome (Table 10). The Hi-C heatmap shows strong interactions within intra-chromosomal region and between paired inter-chromosomes (Fig. 2). Taken together, these results confirm that the allotetraploid oyster genome assembly is of high quality considering its high heterozygosity and repeat content. Alignment of randomly selected genes confirms the presence of both *C. gigas* and *C. angulata* alleles, and the mitochondrial genome is from *C. angulata*, consistent with the known pedigree of the allotetraploid oyster.

**Table 9.** Statistical results of short-read (Illumina) and HiFi-reads (PacBio) alignment.

|  | Illumina | PacBio |
| --- | --- | --- |
| Total reads | 1,052,623,204 | 2,123,642 |
| Mapped reads | 1,043,076,313 | 2,122,608 |
| Mapping rate (%) | 99.09 | 99.95 |
| Average sequencing depth (×) | 74 | 17 |
| Coverage of genome > 1 × (%) | 99.82 | 99.99 |
| Coverage of genome > 5 × (%) | 99.65 | 97.93 |
| Coverage of genome > 10 × (%) | 99.42 | 85.57 |
| Coverage of genome > 20 × (%) | 98.68 | 36.25 |

**Table 10.** The CEGMA and BUSCO assessment of allotetraploid oyster genome assembly.

| Software | Type | Number (percent) |
| --- | --- | --- |
| CEGMA assessment | Number of 458 CEGMA present in assembly | 447 (97.60%) |
|  | Number of 248 highly conserved CEGMA present | 218 (87.90%) |
| BUSCO assessment | Complete and single-copy BUSCOs(S) | 118 (12.37%) |
|  | Complete and duplicated BUSCOs(D) | 820 (85.95%) |
|  | Fragmented BUSCOs(F) | 5 (0.52%) |
|  | Missing BUSCOs(M) | 11 (1.15%) |
|  | Total Lineage BUSCOs | 954 |

## Code availability

No custom code was used during this study for the curation and validation of the dataset. All commands and pipelines used in data processing were executed according to the manual and protocols of the corresponding bioinformatics software.

## Acknowledgements

This work was supported by the National Natural Science Foundation of China (32471687 to A.L.), the Key Research and Development Program of Shandong (ZFJH202309 to G.Z.), the Youth Innovation Promotion Association, Chinese Academy of Sciences (2023215 to A.L.), the Strategic Priority Research Program of the Chinese Academy of Sciences (XDB0730300 to A.L.), the Taishan Scholars Program (tsqn202312267 to A.L.), the Key Research and Development Program of Shandong (2022LZGC015 to L.L.), and the China Agriculture Research System of MOF and MARA (CARS-49 to L.L.).

## Author contributions

G.Z., Xm.G. and L.L. conceived the study. Y.L., X.L., J.F., Xr.G. and Z.X. produced allotetraploids and conducted flow cytometry. A.L., J.Z., M.J.Z., M.S.Z., M.H., J.D., L.W., H.Q. and W.W. collected the samples, extracted the genomic DNA, and conducted sequencing. J.Z. and A.L. performed bioinformatics analysis. A.L., J.Z. and Xm.G. wrote the manuscript. All authors read and approved the final manuscript.

